# Depot-specific roles for sphingosine kinase 1 in conversion of white adipose to thermogenic adipose

**DOI:** 10.1101/2025.10.16.682897

**Authors:** Yolander Valentine, Maryam Jamil, Anna Kovilakath, Ryan D. R. Brown, Jordon Dail, Saaid Farooq, Sarah Spiegel, L Ashley Cowart

## Abstract

**Background:** As of 2023, approximately 100.1 million adults and 14.7 million children in the USA are obese. Many comorbidities develop with obesity, which impairs quality of life and burdens the health care system. Consequently, there is an urgent need for interventions and treatments to reverse obesity and its comorbidities and restore health. Sphingosine Kinase 1 (SphK1), a key enzyme in sphingolipid metabolism, produces sphingosine-1-phosphate (S1P), a bioactive lipid implicated in obesity and metabolic dysfunction. While global deletion of *Sphk1* protects against diet-induced obesity, adipocyte-specific SPHK1 deficiency paradoxically promotes weight gain, glucose intolerance, and adipose inflammation. Given the known role of sphingolipids in adipose thermogenesis, we investigated whether *Sphk1* regulates adipocyte beiging and mitochondrial function.

**Methods:** We assessed thermogenic responses in SphK1-deficient adipocytes and adipocyte-specific *Sphk1* knockout (Ad-SphK1Δ) mice under basal and β3-adrenergic stimulation using CL 316,243. Thermoneutral housing (30°C) and room temperature (23°C) conditions were used to minimize and assess ambient temperature effects on thermogenesis. Molecular, histological, and bioenergetic analyses were conducted across multiple adipose depots.

**Results:** β3-adrenergic stimulation upregulated *Sphk1* expression in mature white adipocytes, while SphK1-deficient adipocytes exhibited enhanced Ucp1 expression, indicating a suppressive role for SphK1 in beiging. In vivo, adipocyte-specific *Sphk1* knockout (Ad-SphK1Δ) mice showed elevated *Ucp1* expression in inguinal and gonadal white adipose tissue (iWAT, gWAT), both basally and after CL 316,243 treatment. These changes were accompanied by depot-specific alterations in adipocyte size and increased adiposity, independent of ambient temperature. Despite similar elevation of thermogenic markers, *Sphk1* deletion had differential effect on mitochondrial function: iWAT showed increased mitochondrial content but reduced complex IV activity and ATP production, whereas gWAT showed reduced mitochondrial abundance without changes in respiration.

**Conclusion:** Our work suggests that *Sphk1* may act as a negative regulator of thermogenic expression and affect mitochondrial function in a depot-specific manner. Loss of *Sphk1* enhances beiging but compromises mitochondrial efficiency, revealing a complex role for the SphK1/S1P axis in adipose plasticity and metabolic regulation. These insights may inform future therapeutic strategies targeting sphingolipid pathways for obesity and metabolic disease.

## Introduction

Obesity and its related metabolic disorders have increased tremendously in the United States and globally, driving an urgent need for novel therapeutic strategies that can effectively mitigate weight gain and its systemic consequences. In recent years, increasing attention has been directed toward understanding the cellular and molecular pathways that govern adipose tissue function, particularly those influencing energy expenditure and thermogenesis.

Among the pathways implicated in obesity, the sphingolipid signaling axis has emerged as a critical regulator of metabolic homeostasis. Sphingosine-1-phosphate (S1P), a bioactive lipid produced by sphingosine kinase 1 (SPHK1), is elevated in the plasma of obese individuals and plays pleiotropic roles in inflammation, insulin resistance, and lipid metabolism [1]. Consequently, both SPHK1 and its product S1P are increasingly recognized as important contributors to obesity and its associated metabolic disturbances [2-4]. While global deletion of *Sphk1* has been shown to protect against diet-induced obesity and systemic inflammation in mice [5], our laboratory has demonstrated that adipocyte-specific deletion of *Sphk1* paradoxically worsens metabolic outcomes [6]. Mice lacking *Sphk1* in adipocytes exhibit increased weight gain, adipocyte hypertrophy, glucose intolerance, and hepatic steatosis when exposed to a high-fat diet [6], suggesting a context- and cell type-specific role for the SphK1/S1P axis in metabolic regulation.

Thermogenesis—the physiological process of heat production—plays a critical role in maintaining energy balance by promoting energy expenditure. It serves as an effective mechanism for dissipating excess energy and has been shown to counteract diet-induced obesity in various preclinical models [7-10]. This process is primarily mediated by specialized adipocytes, brown and beige fat cells, which possess high mitochondrial content and express uncoupling protein 1 (UCP1), a key driver of non-shivering thermogenesis. β3-adrenergic receptor (β3AR) activation is a well-characterized stimulus for thermogenesis, promoting UCP1 expression and mitochondrial uncoupling in both brown and beige adipocytes. While the sphingolipid metabolite ceramide has been implicated in modulating adipocyte differentiation, mitochondrial function, and thermogenesis [11-15], the specific contributions of adipocyte SPHK1 and S1P to thermogenic regulation remain undefined.

To address this gap, we hypothesized that adipocyte *Sphk1* regulates thermogenic capacity in white adipose tissue, particularly in response to β3AR signaling. Using a conditional adipocyte-specific *Sphk1* knockout (Ad-SphK1Δ) mouse model, we examined the role of *Sphk1* in thermogenesis under both basal and pharmacologically stimulated conditions. We assessed the effects of *Sphk1* deletion on core body temperature, UCP1 expression, adipocyte morphology, and mitochondrial function, with and without treatment using CL 316,243, a selective β3AR agonist. This experimental framework allowed us to determine whether *Sphk1* contributes to, or constrains, the thermogenic remodeling of adipose tissue and how it intersects with mitochondrial energetics.

By clarifying the role of the SphK1/S1P axis in adipocyte thermogenic programming, this study aims to fill a critical knowledge gap in the field of sphingolipid biology and metabolic regulation. A deeper understanding of how *Sphk1* modulates mitochondrial health and energy expenditure may ultimately lead to novel therapeutic strategies targeting adipose tissue plasticity for the treatment of obesity and its comorbidities.

## Results

### *Sphk1* expression is induced by β3-adrenergic receptor activation in mature white adipocytes

We previously demonstrated that adipocyte-specific deletion of sphingosine kinase 1 (*Sphk1*) promotes weight gain in mice fed a high-fat diet, accompanied by adipocyte hypertrophy, impaired glucose tolerance, and reduced lipolysis [6]. These findings led us to investigate whether *Sphk1* plays a role in adipose tissue beiging.

To explore this, stromal vascular cells (SVC) were isolated from control and global *Sphk1* knockout (*Sphk1*^*-/-*^) mice and differentiated into mature white adipocytes. Cells were then treated with CL 316,243 (CL), a selective β3-adrenergic receptor agonist known to induce adipose beiging. In control adipocytes, CL treatment significantly upregulated *Sphk1* mRNA expression (Fig. 1A), indicating that *Sphk1* is induced downstream of β3-adrenergic signaling, consistent with previous reports [16, 17]. To determine whether loss of *Sphk1* was compensated by *Sphk2*, we measured *Sphk2* mRNA expression. Although CL robustly reduced *Sphk2* expression, no significant changes were observed in *Sphk1*^*-/-*^ adipocytes, irrespective of the treatment (Fig. 1C).

**Figure 1:**
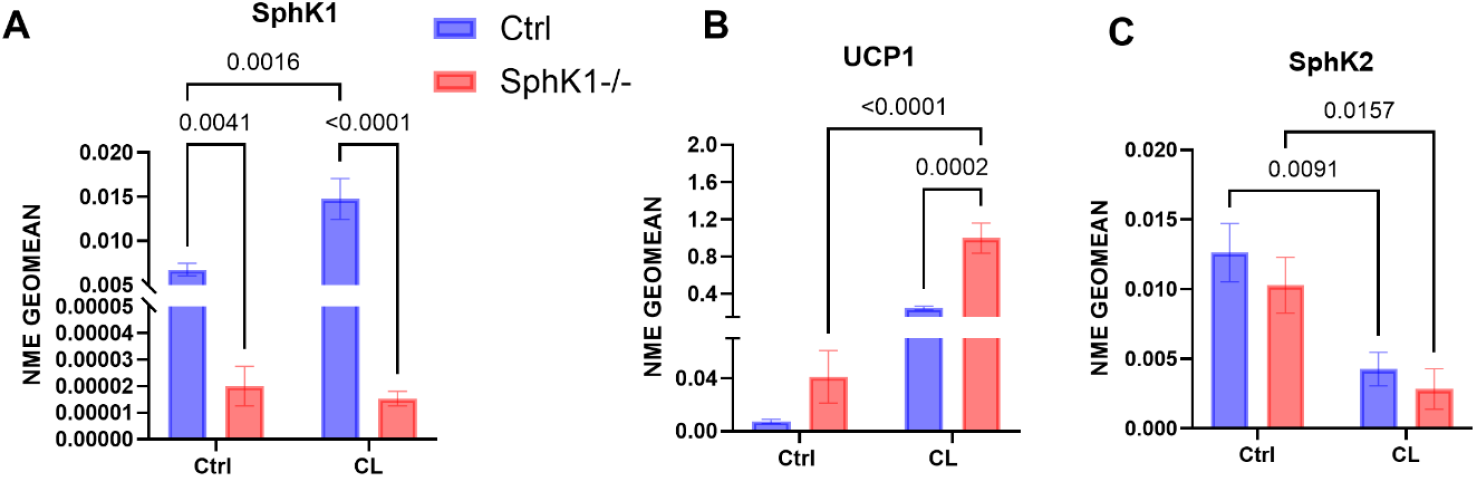
*Sphk1* expression is induced by β3-adrenergic receptor activation in mature white adipocytes. Stromal vascular cells from inguinal white adipocytes from control and *Sphk1*^*-/-*^ mice were isolated and differentiated into mature white adipocytes and treated with 2 µM CL316,243 (CL) for 16 hours. mRNA expression of *Sphk1* (A), *Ucp1 (B)*, and *Sphk2 (C)* (N=3/group). Data expressed as mean ±SEM. Statistical analysis was performed using two-way ANOVA; p-values are indicated.

We next assessed thermogenic gene expression by measuring *Ucp1*, a canonical marker of adipocyte beiging. As expected, CL treatment induced *Ucp1* expression in both genotypes, but the increase was significantly greater in *Sphk1* deleted adipocytes compared to controls (Fig. 1B). This enhanced thermogenic response in the absence of *Sphk1* suggests that it may negatively regulate adipocyte beiging.

### *Sphk1* depletion enhances the beiging response to β3-adrenergic receptor stimulation in vivo

To minimize basal thermogenic gene expression, *Sphk1*^fl/fl^ Adipoq-Cre^+^ (Ad-SphK1Δ) mice and their *Sphk1*^fl/fl^ littermate controls were housed at thermoneutrality (30°C) for 2 weeks prior to treatment. This eliminated baseline differences in UCP1 expression caused by chronic cold stress at standard housing temperatures (∼23°C). Following this acclimation period, mice were administered daily intraperitoneal injections of either CL 316,243 (1 mg/kg) or saline for 10 days while remaining at thermoneutrality (Fig. 2A).

**Figure 2:**
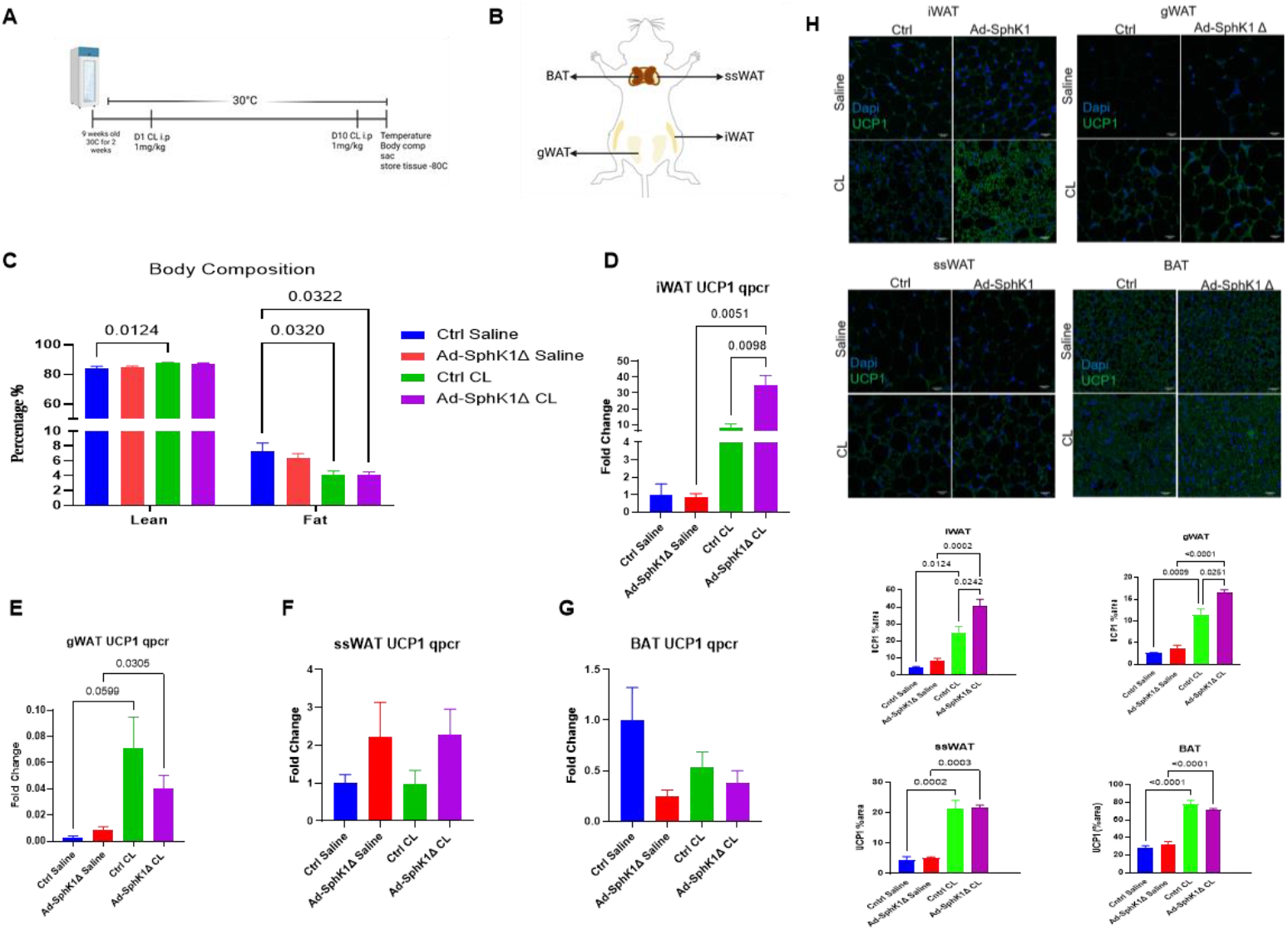
*Sphk1* depletion enhances adipocyte response to β3 adrenergic receptor stimulation. Adipose beiging was induced in control and Ad-SphK1Δ mice housed at thermoneutrality (30ºC) by intraperitoneal injection of the β3AR selective agonist CL316,243 (1 mg/kg). Adipose tissues were collected and analyzed for induction of thermogenic genes. Representative schematic of **A**. thermoneutrality experiment, and **B**. location of the four adipose depots. **C**. Whole body composition was analyzed using Burker’s minispec analyzer (N=10/group). UCP1 expression of **D**. iWAT, **E**. gWAT, **F**. ssWAT, and **G**. BAT (N=6-8/group). **H**. Representative 40x (scale bar = 20 µm) magnified adipose depots for UCP1 staining and quantification (N=3, 4 biological replicates/group, N=120-150 adipocytes/group). Data expressed as mean ±SEM. C. Two way ANOVA with FDR-adjusted p values shown. D. to H. One way ANOVA with FDR-adjusted p values shown.

To assess depot-specific effects of *Sphk1* on adipose beiging, four adipose depots were collected: inguinal WAT (iWAT); gonadal WAT (gWAT); subscapular WAT (ssWAT); and brown adipose tissue (BAT) (Fig. 2B). Although, there were no major changes in body weight, CL treatment as expected reduced fat mass (Fig. 2C). Ad-SphK1Δ mice displayed slightly lower body temperatures compared to controls in both saline- and CL-treated groups, though the differences were not statistically significant (Suppl. Fig. 1A). Body weight and total fat mass were also comparable between genotypes (Suppl. Fig. 1B). While depot weights did not differ between genotypes, CL treatment reduced the mass of all three white adipose depots (iWAT, gWAT, and ssWAT), but not BAT (Suppl. Fig. 1C). These differences were not unsurprising because continuous housing at thermoneutrality reduces basal energy expenditure to maintain body temperature.

At the molecular level, CL significantly induced *Ucp1* mRNA expression in iWAT (Fig. 2D) and gWAT (Fig. 2E), with greater induction observed in Ad-SphK1Δ mice compared to controls in iWAT. No genotype-dependent differences in *Ucp1* mRNA expression were observed in ssWAT (Fig. 2F) or BAT (Fig. 2G). These findings were corroborated by UCP1 immunofluorescence staining, which confirmed significantly enhanced protein expression in iWAT from CL-treated Ad-SphK1Δ mice (Fig. 2H).

### Adipocyte-specific *Sphk1* deletion has depot-specific effects on adipocyte hypertrophy

To assess whether differences in UCP1 expression were accompanied by morphological changes, we quantified adipocyte size across various fat depots. At baseline, Ad-SphK1Δ mice exhibited smaller adipocytes in iWAT, ssWAT, and BAT, but larger adipocytes in gWAT compared to *Sphk1*^fl/fl^ controls (Fig. 3A–D). Following CL treatment, all groups significant showed adipocyte hypotrophy. However, in Ad-SphK1Δ mice, CL treatment paradoxically increased adipocyte size in iWAT relative to CL-treated controls (Fig. 3A). In contrast, gWAT adipocyte size in Ad-SphK1Δ mice was significantly reduced by CL, opposite to the pattern observed in control untreated mice (Fig. 3B). These findings highlight a depot-specific role for *Sphk1* in regulating adipocyte morphology and suggest that *Sphk1* depletion disrupts the typical relationship between adipocyte size and UCP1 expression in thermogenic of visceral fat depots.

**Figure 3:**
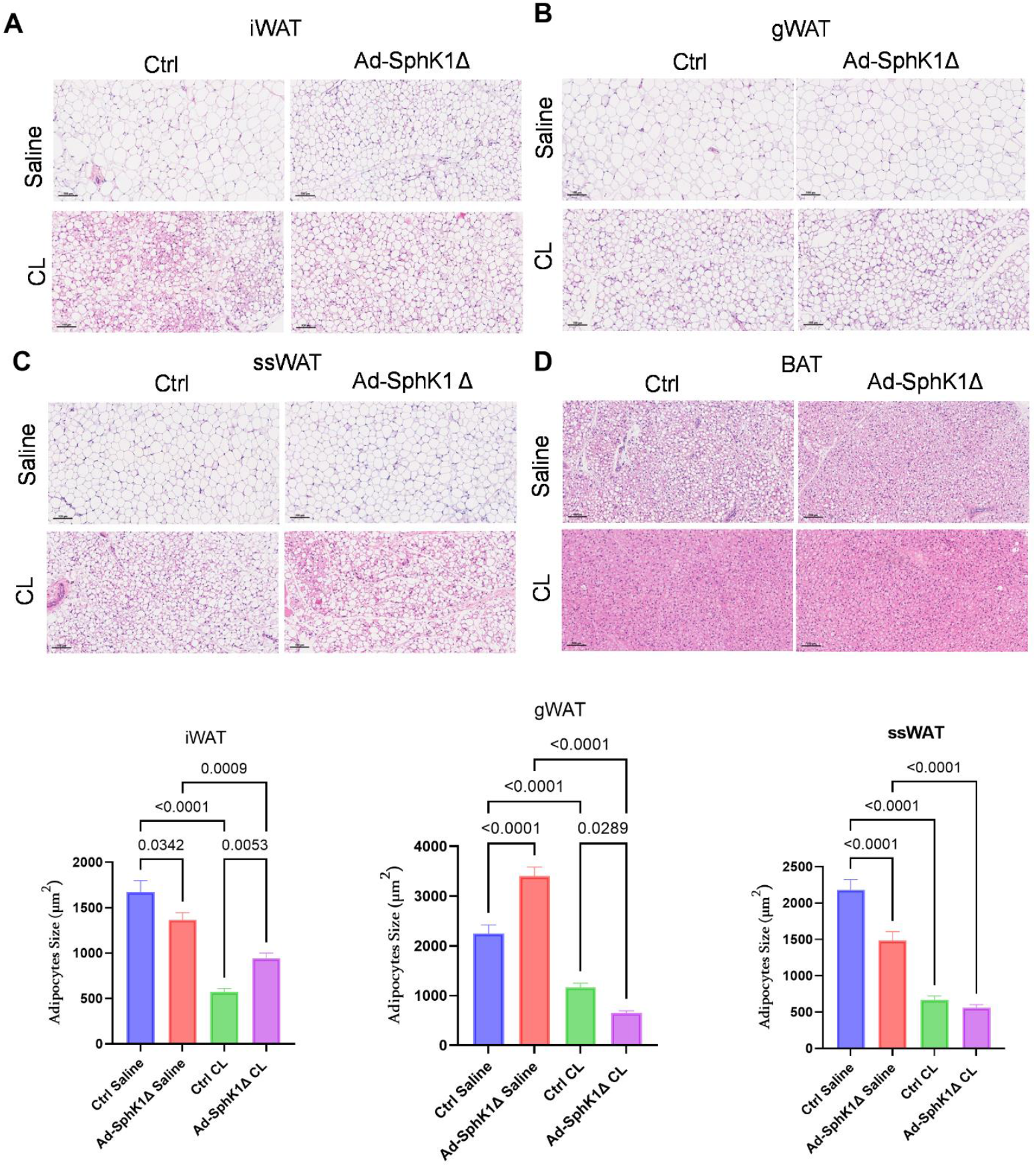
*Sphk1* has a depot specific role in adipose hypertrophy. Adipocyte size was quantified using the ImageJ adiposoft plugin on hematoxylin and eosin stained adipose sections from control and Ad-SphK1Δ mice housed at thermoneutrality treated with saline or CL. Representative 10x (scale bar = 100 µm) magnified adipose depots of **A**. iWAT, **B**. gWAT, **C**. ssWAT, and **D**. BAT (N=3 biological replicates/group, N=120-150 adipocytes/group). Data expressed as mean ±SEM. A. to C. One way ANOVA with FDR-adjusted p values shown.

### *Sphk1* depletion increases adiposity and alters thermogenic programming

Given the depot-specific differences observed in response to CL-induced beiging at thermoneutrality, we next assessed whether these alterations were present under basal conditions. Notably, Ad-SphK1Δ mice displayed increased body weight and slightly elevated core body temperature compared to control littermates (Fig. 4A–B). Adipose depot weights were also significantly higher in Ad-SphK1Δ mice, particularly in iWAT and gWAT (Fig. 4C).

**Figure 4:**
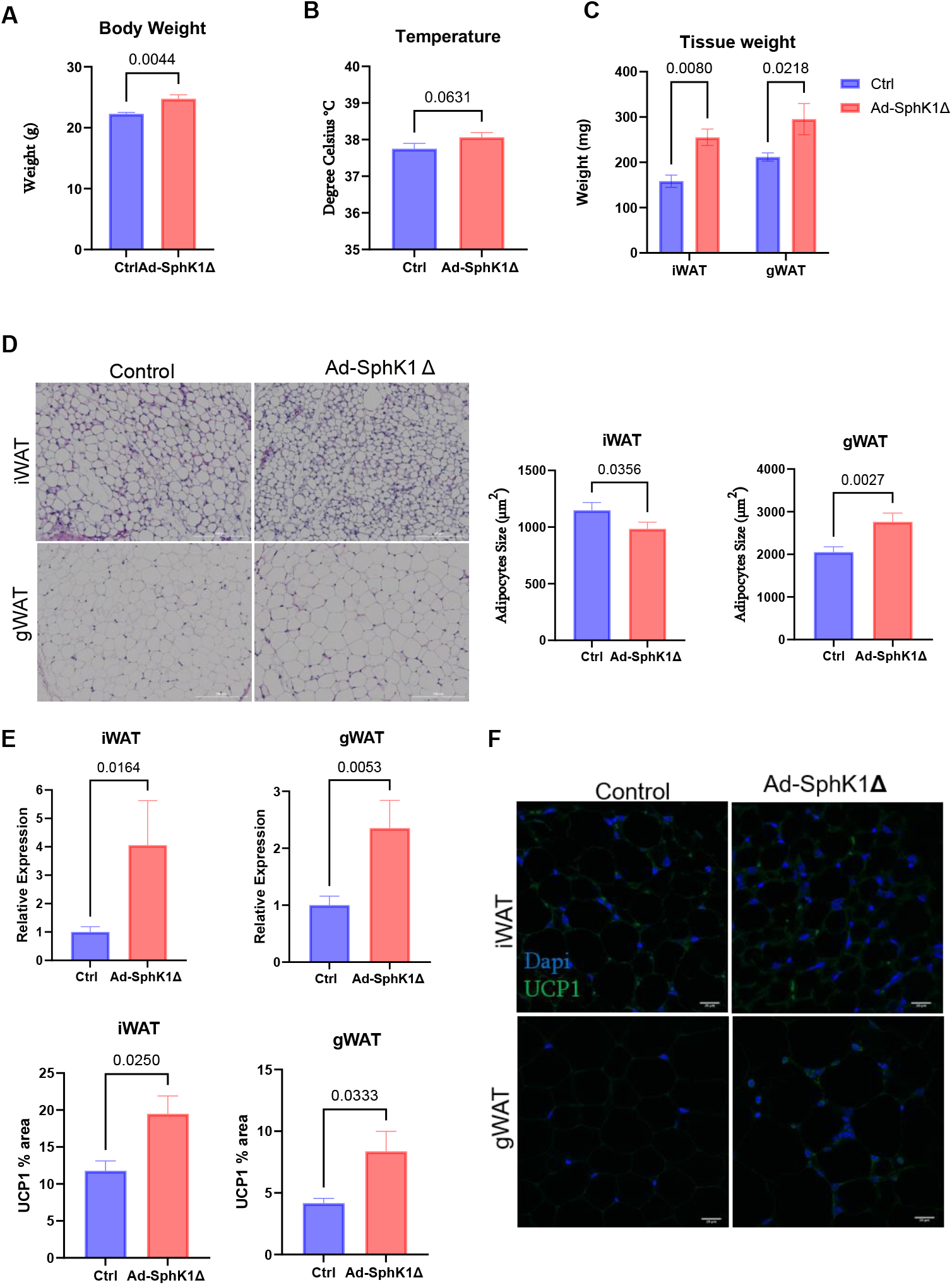
*Sphk1* depletion increased adiposity. Control and Ad-SphK1Δ mice were housed at 23ºC. Measurement of **A**. body weight (N=6/group), **B**. body temperature (N=14-16/group), and **C**. iWAT and gWAT depots (N=6/group). **D**. Adipocyte size was quantified using hematoxylin and eosin stained adipose sections (N=3 biological replicates/group, N=120-150 adipocytes/group). **E**. mRNA expression of *Ucp1* in iWAT and gWAT depots (N=6-9/group). **F**. Representative 40x (scale bar = 20 µm) magnified adipose depots for UCP1 (N=3/group). Data expressed as mean ±SEM. A., B., D. to F. Multiple unpaired Student’s t-test with p values shown. C.

Histological analysis revealed a pattern consistent with thermoneutral findings, that Ad-SphK1Δ mice exhibited adipocyte hypotrophy in iWAT and hypertrophy in gWAT relative to controls (Fig. 4D). To further assess the thermogenic phenotype, we measured UCP1 expression. *Sphk1* depletion led to a marked upregulation of UCP1 in both iWAT and gWAT (Fig. 4E–F), suggesting enhanced beiging in these depots.

Collectively, these results indicate that loss of *Sphk1* promotes adiposity while simultaneously enhancing thermogenic gene expression in a depot-specific manner, independent of ambient temperature. Based on these findings, all subsequent experiments were conducted at room temperature (23°C).

### *Sphk1* deletion reduces mitochondria function in iWAT and gWAT

UCP1 is essential for mitochondrial health as its main role is to dissipate the proton gradient generated from the electron transport chain (ETC). When UCP1 is induced, adipocytes must increase their oxygen consumption rates to maintain the proton gradient for ATP production[18-20]. Therefore, we investigated the role *Sphk1* plays in regulating mitochondrial number and function.

To determine the relative mitochondrial content, we employed two distinct techniques. The first technique involved assessing mitochondrial DNA copy number (Fig. 5A-B), while the second technique utilized flow cytometry to analyze isolated mitochondria stained with MitoTracker Green (Fig. 5C-D). Depletion of *Sphk1* in iWAT led to an increase in mitochondrial content (Fig. 5A,C). In contrast, gWAT from Ad-SphK1Δ mice showed a significant reduction in mitochondrial abundance compared to controls (Fig. 5B, D).

**Figure 5:**
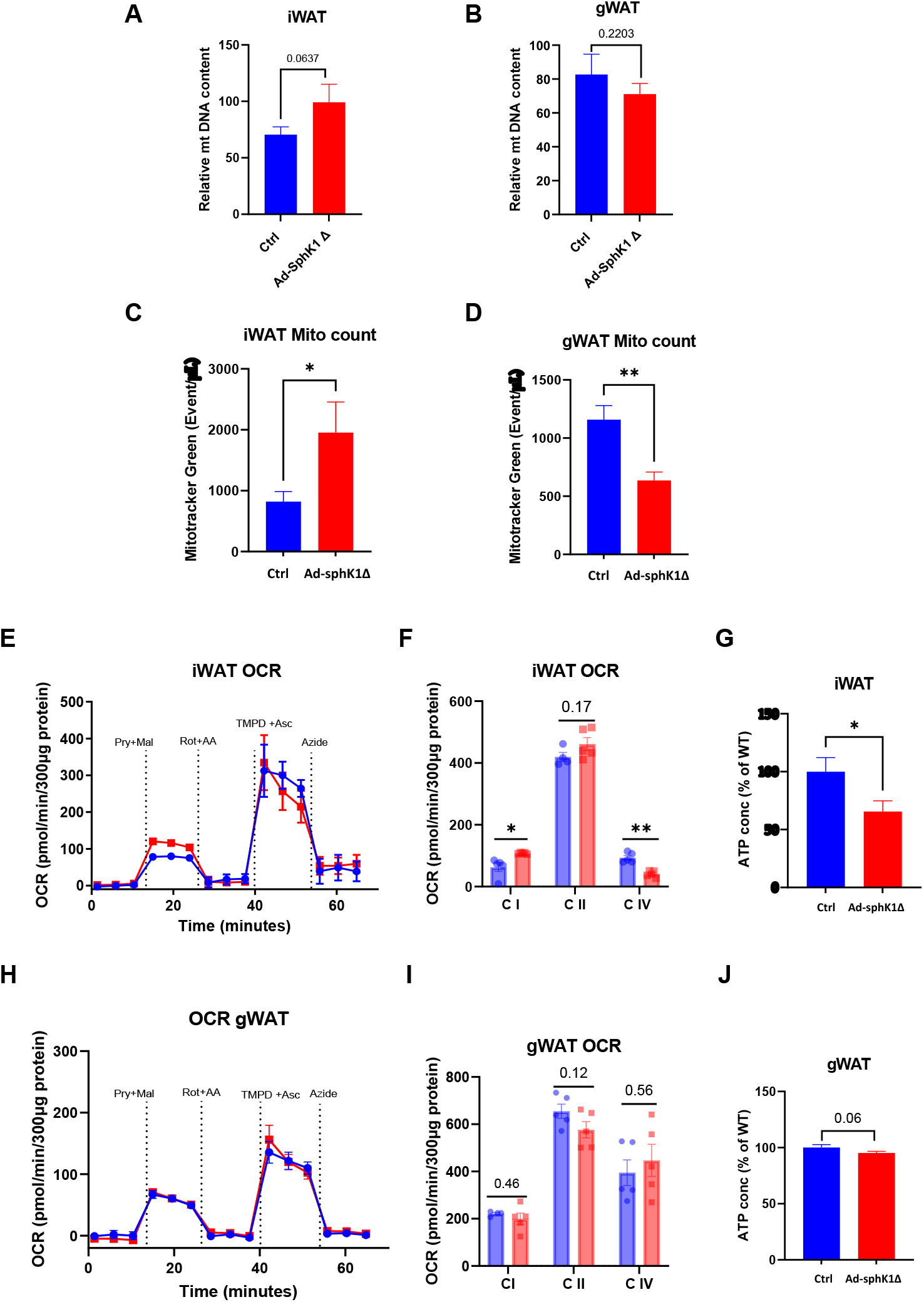
*Sphk1* deletion reduces mitochondria function in iWAT and gWAT. Mitochondria were isolated from control and Ad-SphK1Δ mice and analyzed for mitochondria health and function. Mitochondrial DNA was isolated and expression was quantified by comparing the ratio of mitochondrial ND1, and Mt-co2 to nuclear 18s rna and HK2 in **A**. iWAT, and **B**. gWAT (N=3-4/group). Flow cytometry on 100 µg isolated mitochondria stained with MitoTracker Green for relative mitochondria count in **C**. iWAT, and **D**. gWAT (N=3/group). Seahorse analysis was performed for oxygen consumption rate of mitochondrial electron transport chain complexes on **E**. iWAT, followed by **F**. quantification, and **G**. ATP analysis. Seahorse analysis was performed on **H**. gWAT, followed by **I**. quantification, and **J**. ATP analysis. Data expressed as mean ±SEM. A. to D., G., J. one-tailed t-test with p values shown. F. and I. One way ANOVA with p values shown.

To evaluate mitochondrial function, we performed Seahorse respirometry to measure ETC complex-specific oxygen consumption rates (OCR) [21, 22]. In iWAT, *Sphk1* deletion enhanced complex I OCR but reduced complex IV OCR (Fig. 5E–F). Additionally, total ATP production was significantly decreased in iWAT from Ad-SphK1Δ mice (Fig. 5G). In gWAT, however, *Sphk1* loss did not significantly affect OCR or ATP production (Fig. 5H–J).

Together, these data demonstrate that *Sphk1* plays a critical, depot-specific role in maintaining mitochondrial integrity and function in adipose tissue, particularly in thermogenic iWAT.

## Discussion

Our study reveals a previously unrecognized role for *Sphk1* in regulating thermogenic programming in a depot-specific manner. We demonstrate that *Sphk1* modulates UCP1 expression both under basal conditions and in response to β3-adrenergic stimulation. Interestingly, although deletion of *Sphk1* in adipocytes resulted in elevated UCP1 expression in iWAT and gWAT, this did not translate into the expected metabolic phenotype—namely, reduced body weight, adipose mass, or adipocyte size. Instead, Ad-SphK1Δ mice exhibited increased weight gain and adiposity despite enhanced thermogenic gene expression.

This paradox was especially evident under both room temperature and thermoneutral housing. At thermoneutrality, where baseline sympathetic tone is minimized, Ad-SphK1Δ mice failed to mount a thermogenic response to CL 316,243 in terms of body temperature elevation—despite exhibiting robust UCP1 upregulation, particularly in iWAT. Notably, iWAT adipocytes in these mice were paradoxically larger following CL treatment, indicating that increased UCP1 expression did not drive the expected lipid mobilization and adipocyte hypotrophy. Supporting this, our data show that Ad-SphK1Δ mice displayed impaired lipolytic responses to β3-adrenergic stimulation, as reflected by the lack of circulating free fatty acid elevation—a key energy source for UCP1-mediated thermogenesis (Suppl. Fig. 2) [23-25].

The disconnect between increased UCP1 expression and thermogenic efficacy appears to be linked to mitochondrial dysfunction. In iWAT from Ad-SphK1Δ mice, mitochondrial numbers were increased, yet functional analysis revealed reduced complex IV oxygen consumption, diminished mitochondrial membrane potential, and decreased ATP production. High UCP1 levels may reduce the proton gradient and make it less available for complex IV. This can reduce the rate of electron transfer through complex IV and potentially lower oxygen consumption due to decreased production of ATP.These findings suggest a compensatory response to mitochondrial inefficiency, wherein increased mitochondrial biogenesis and UCP1 expression are unable to overcome the functional deficits caused by SPHK1 loss. Thus, *Sphk1* appears to be essential not only for thermogenic gene regulation but also for maintaining mitochondrial integrity and bioenergetic output.

Our results also point to the predominant role of brown adipose tissue in systemic temperature regulation. At room temperature, both body temperature and BAT UCP1 expression were elevated in Ad-SphK1Δ mice. However, when housed at thermoneutrality, where BAT thermogenesis is largely suppressed, body temperature in the mutant mice declined despite elevated UCP1 in white depots. This supports the notion that UCP1 expression in BAT, rather than in beige depots, is the primary driver of core body temperature maintenance. It also suggests that *Sphk1* may play a role in thermoregulation through effects in BAT, though future studies are needed to explore this directly.

Together, these findings identify *Sphk1* as a regulator of adipose thermogenic capacity, with distinct and depot-specific effects on UCP1 expression, adipocyte morphology, and mitochondrial function. Importantly, our data suggests that elevated UCP1 expression is not synonymous with enhanced thermogenesis—particularly in the absence of mitochondrial integrity. The SphK1/S1P axis emerges as a potential new signaling node required for coupling thermogenic transcriptional programs with mitochondrial function and energy dissipation.

In summary, this study provides new insight into the role of sphingolipid signaling in adipose tissue biology, revealing that *Sphk1* can contribute to regulation of thermogenesis and mitochondrial homeostasis. These findings underscore the complexity of adipose tissue regulation and suggest that targeting SphK1/S1P signaling may offer novel therapeutic avenues for obesity and metabolic disorders.

## Methods

### Animal model

All animal experiments conformed to the protocols approved by the institutional animal care and use committee. Mice were housed in the animal facility Virginia Commonwealth University (VCU). Food and water were provided ad libitum. Animals were maintained on a 12h:12h light:dark cycle. Tissues were collected accordingly as fresh fixed in 10% neutral buffered formalin or fresh snap frozen in liquid nitrogen and stored at -80°C until further processing. All experiments were approved by the Virginia Commonwealth University Institutional Animal Care and Use Committee.

*In Vivo* Beiging *Sphk1* Fl/Fl AdipoCre- (*Sphk1*^fl/fl^) and *Sphk1* Fl/Fl Adipo Cre+ (Ad-SphK1Δ) mice 10 weeks old were housed at 30°C for two weeks. After which these mice were injected with CL316,243 (TOCRIS 1499) 1mg/kg or saline control daily for ten days, while continued being housed at 30°C. Mice were euthanized one day post injection and adipose collected and snap frozen in liquid nitrogen and stored at -80°C. Tissue for histology were fixed in 10% neutral buffered formalin for 24 hours and then stored in 70% ethanol.

### Quantitative PCR

RNA isolation was done as previously described [6] using Qiagen (74536) total RNA isolation kit. For tissue lysate, 100 µL of homogenate was added to 400 µL of TRIzol and RNA isolation was done as described above. Complementary DNA (cDNA) was made using BioRad iScript™ cDNA Synthesis (1708890). Quantitative PCR (qPCR) was performed with BioRad ssoAdvanced Universal SYBR® Green Supermix (1725274). Geomean of multiple housekeeping genes was used to normalize Cq values.

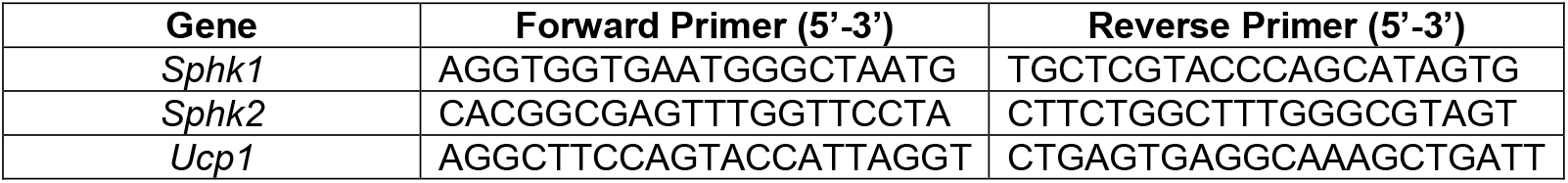

### Mitochondrial DNA copy number

For DNA isolation from adipose tissue 10 mg of tissue was homogenized at 20 s for 15 Hz in Buffer ATL. Steps for Qiagen, DNeasy® Blood & Tissue kit, 69504 was used. For each sample, 6 ng of DNA was added to each well of a qPCR plate. The Cq values of the mitochondrial gene MT-CO2 were determined, and the Cq values of the nuclear gene 18s RNA were also measured. To determine the mitochondrial DNA copy number, the following formula was used. (a) ΔCT = (nucDNA CT − mtDNA CT). (b) Relative mitochondrial DNA content = 2 × 2ΔCT.

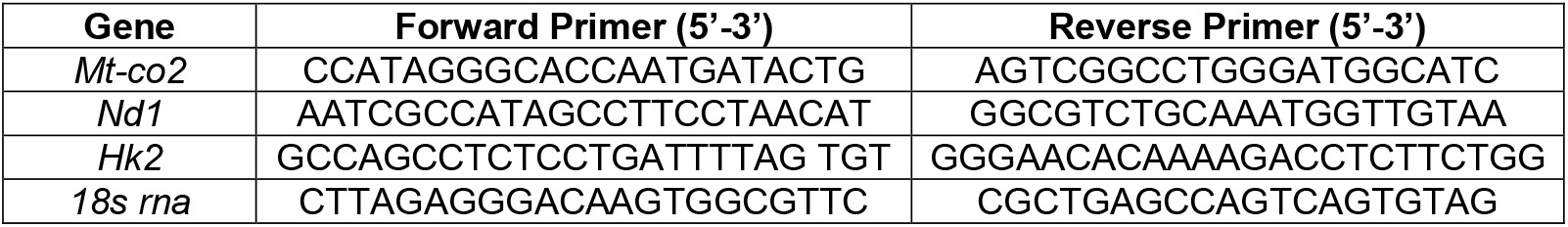

### Seahorse extracellular flux

Seahorse for adipose depots was done as previously described [21, 22]. Briefly, tissue was homogenized in glass pestle, lysates were then centrifuged at 3000g for 10 minutes at 4°C. BCA was done to determine protein concentration and 125ug of protein was loaded on V7 seahorse plates. Tissue homogenate and cell for mitochondrial complex seahorse plates in 1X MAS buffer (Mannitol 220Mm, sucrose 70mM, KH2P04 5mM, HEPES 2mM, EGTA 1mM, fatty free BSA 0.2% (W/V)). Seahorse OCR for mitochondrial complexes was adopted from [21, 22]. For complex I and IV OCR pyruvate 10mM and malate 10mM port A; Rotenone 2µM and Antimycin A 4 µM port B; TMPD 0.25 mM and Ascorbic acid 0.5mM port C; and Sodium Azide 50mM port D. For complexes II and IV OCR succinate 5mM and Rotenone 2µM port A; Rotenone 2µM and Antimycin A 4 µM port B; TMPD 0.25 mM and Ascorbic acid 0.5mM port C; and Sodium Azide 50mM port D. The calculations for each complex were as follows Complex I maximal respiration OCR_pry+mal_ – OCR_ROT+AA_, complex II maximal respiration OCR_succ+ROT_ – OCR_ROT/AA_, complex IV OCR_TMPD+ASC_-OCR_Azide_.

### Mitochondria isolation

Mitochondria was isolated as previously described [26] from freshly isolated adipose tissue. Adipose was washed in ice cold PBS. Adipose tissue was finely minced with scissors then homogenized in Media 1 (250mM sucrose, 10mM Hepes, 1mM EGTA), homogenate was centrifuged at 8500g for 10 minutes at 4^°^C. The pellet was resuspended in ice cold Media 1 and centrifuged at 800g for 10 minutes. The resulting supernatant was centrifuged at 8500g for 10 minutes. The pellet was resuspended in ice cold Media 2 (100mM KCl, 20mM Hepes, 1mM EDTA, 0.6% fatty acid free BSA, pH 7.2) and centrifuged for 10 minutes at 8500g, final mitochondrial pellet was suspended in Media 2 and BCA was used to determine concentration. Isolated mitochondria were stained for flow cytometry, used for Complex IV immunoprecipitation, and seahorse.

### Flow cytometry

Isolated mitochondria were stained with 80nM MitoTracker Green (M7514) and 80nM Mito Tracker Orange (M7510) in 1X MAS buffer for 15 minutes at 4C in the dark. Mitochondria were washed by centrifuging at 17000g for 2 minutes and then resuspended in 1X MAS buffer (Mannitol 220Mm, sucrose 70mM, KH2P04 5mM, HEPES 2mM, EGTA 1mM, fatty free BSA 0.2% (W/V)). Multicolor data acquisition was performed using Cytek Aurora (Cytek) as previous described [27, 28]. Data are analyzed using FlowJo, version 10.8.0. Services in support of the research project were provided by the VCU Massey Cancer Center Flow Cytometry Shared Resource supported, in part, with funding from NIH-NCI Cancer Center Support Grant P30 CA016059. Mitochondria count was normalized to Event/uL by the volume ran. Cytek instrument was set to log mode and voltages set as follows: FSC: 450; SSC 250. Events were collected in FSC-A (area) mode, FSC-W (width), FSC-H (height). 200,000 events were collected.

### ATP assay

Mitochondria was isolated as previously described [26] from freshly isolated adipose tissue. Adipose was washed in ice cold PBS. Adipose tissue was finely minced with scissors then homogenized in Media 1 (250mM sucrose, 10mM Hepes, 1mM EGTA), homogenate was centrifuged at 8500g for 10 minutes at 4^°^C. The pellet was resuspended in ice cold Media 1 and centrifuged at 800g for 10 minutes. The resulting supernatant was centrifuged at 8500g for 10 minutes. The pellet was resuspended in ice cold Media 2 (100mM KCl, 20mM Hepes, 1mM EDTA, 0.6% fatty acid free BSA, pH 7.2) and centrifuged for 10 minutes at 8500g, final mitochondrial pellet was suspended in Media 2 and BCA was used to determine concentration. 100ug of frozen isolated mitochondria was used. Manufactor’s instruction was followed for ATP Assay kit ab83355.

### Immunofluorescence staining and confocal microscopy

Paraffin sections of adipose were dewaxed and rehydrated by submerging for 5 minutes, twice in Histoclear, 90% Histoclear/10% ethanol, twice in 100% ethanol, twice in 95% ethanol, twice in 80% ethanol, twice in 70% ethanol before submerging twice in distilled water for 20 minutes. HIER antigen-retrieval buffer (#208572, Abcam) was heated to 98°C and slides immediately submerged in solution for 20 minutes before being washed in PBS for 5 minutes, 3 times. Slides were placed in in ice-cold methanol for 15 minutes at -20°C for fixation and subsequently washed in PBS for 10 minutes, 3 times. Tissue sections were then covered with a layer of PBS containing 5% normal goat serum (#31872, ThermoFisher), 1% BSA and 0.4% Triton X-100 (buffer A) for 1 hour at room temperature to simultaneously block and permeabilise. Sections were incubated overnight at 4°C with anti-UCP1 (1:200 in buffer A) (#10983, Abcam) and then washed in PBS for 10 minutes, 3 times before being incubated for 1 hour with anti-rabbit Alexa Fluor™ 488 (1:200 in buffer A) (#A11008, ThermoFisher) at room temperature. Slides were then washed in PBS for 5 minutes, 3 times and mounted with VECTASHIELD Vibrance antifade mounting medium containing DAPI (#H-1800, Vector Laboratories).

16-bit Images were captured using an inverted Zeiss LSM880 confocal microscope. A 405 nm laser diode and a 488 nm argon-ion laser were sequentially used to excite DAPI and Alexa Fluor™ 488, respectively. The confocal pinhole was adjusted to 1 Airy unit for an emission of 520 nm, detecting emission of DAPI at 410-470 nm and Alexa Fluor™ 488 at 490-550 nm. Analysis of 3-5 biological replicates per group was done using ImageJ to measure the % of positive stained area for UCP1 after thresholding all images to 2500-65535.

### Statistical analysis

All values are reported as mean ± SEM (standard error of the mean). When comparing two groups, pairwise comparisons of normally distributed datasets were conducted using a Welch’s corrected two-tailed Student t-test. For comparisons involving multiple groups, either a one-way ANOVA or a two-way ANOVA with the appropriate correction test was performed. These statistical analyses were carried out using GraphPad Prism 9 software.

## Acknowledgements

We thank Dr. Francesco Celi and Bin Ni for their expertise in adipose thermogenesis and guidance throughout this project. We also thank you for allowing us to use your equipment, like the body composition analyzer. We thank Dr. Johana Lambert for her contributed knowledge of adipose biology. We thank Dr. Rebecca Martin, Madison Isbell and the VCU Flow Core staff for the guidance and assistance with the flow cytometry experiments.

This study and its personnel were supported in part by grants from the National Institutes of Health 5R01HL151243-04, 1R21AA029518-01A1, and Veterans’ Affairs 5I01BX000200-13 to L.A.C., F31HL156529, T32HL149645 to AK, 1F31DK129028 to YV, and R35GM152058 to S.S.. Services and products in support of the research project were generated by the following Lipidomics and Metabolomics Shared Resource, supported, in part, with funding from NIH-NCI Cancer Center Support Grant P30 CA016059. Virginia Commonwealth University Flow Cytometry Shared Resource, supported, in part, with funding from NIH-NCI Cancer Center Support Grant P30 CA016059. VCU Tissue and Data Acquisition and Analysis Core (TDAAC) Facility, supported, in part, with the funding from NIH-NCI Cancer Center Core Support Grant P30 CA016059, as well as through the Dept. of Pathology, School of Medicine, and Massey Cancer Center of Virginia Commonwealth University

## Authors Contribution

LAC and SS conceived and directed the project. YV designed and carried out experiments. MJ and AK assisted with sample collection. RB performed IF staining. JD and SF assisted with qPCR. YV, MJ, and AK wrote the original draft. LAC, and SS edited the manuscript.

## Supplemental Figures and Figure Legends

**Supplementary Figure 1:**
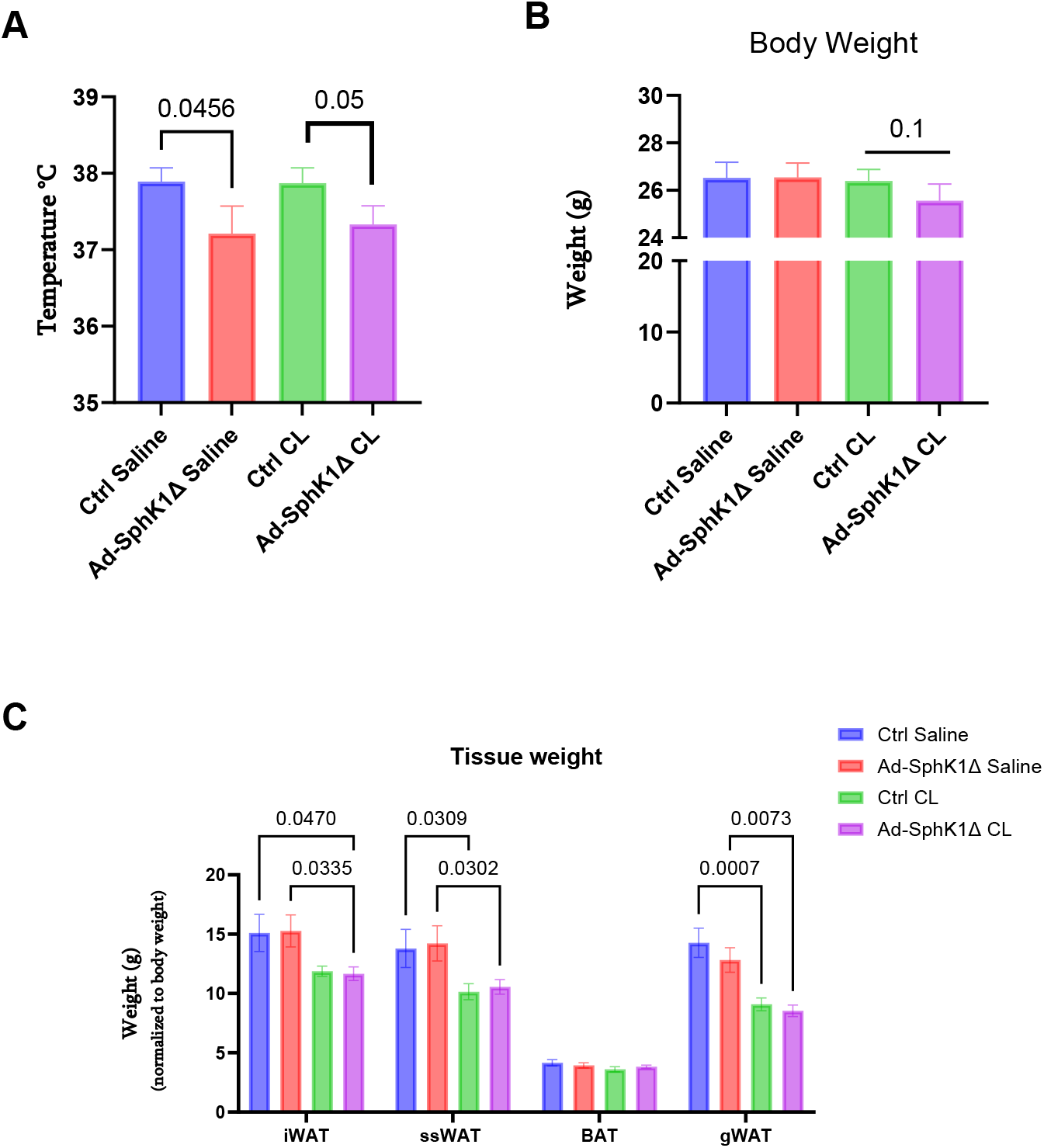
Ad-SphK1Δ mice housed at thermoneutrality show reduced body temperature, body weight, and fat mass compared to control mice. **A**. Body temperature, **B**. body weight, and **C**. Adipose tissue weights (iWAT, gWAT, ssWAT, BAT) after CL treatment in Cntrl and Ad-SphK1Δ mice(N=10-13/group). Data expressed as mean ±SEM. A. and B. Unpaired Student’s t-test with p values shown. C. Two way ANOVA with FDR-adjusted p values shown.

**Supplementary Figure 2:**
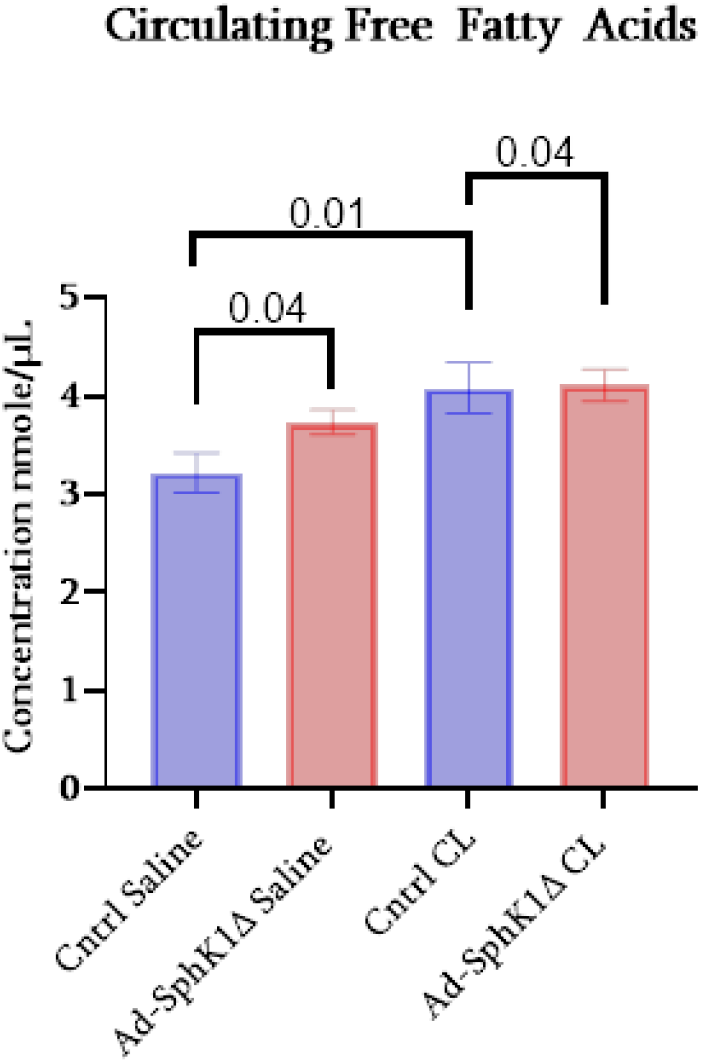
Decrease in circulating Free Fatty Acids in Ad-SphK1Δ mice with CL treatment. **A**. CL injection induced circulating free fatty acids in control mice but not in Ad-SphK1Δ mice(N=5-8/group). Data expressed as mean ±SEM. Unpaired Student’s t-test with p values shown.

## References

1. Kowalski, G.M., et al., Plasma sphingosine-1-phosphate is elevated in obesity. PLoS One, 2013. 8(9): p. e72449.

2. Valentine, Y. and L.A. Cowart, Sphingolipids in Adipose: Kin or Foe? Adv Exp Med Biol, 2022. 1372: p. 15–29.

3. Green, C.D., et al., Sphingolipids in metabolic disease: The good, the bad, and the unknown. Cell Metab, 2021. 33(7): p. 1293–1306.

4. Lambert, J.M., A.K. Anderson, and L.A. Cowart, Sphingolipids in adipose tissue: What’s tipping the scale? Adv Biol Regul, 2018. 70: p. 19–30.

5. Wang, J., et al., Sphingosine kinase 1 regulates adipose proinflammatory responses and insulin resistance. Am J Physiol Endocrinol Metab, 2014. 306(7): p. E756–68.

6. Anderson, A.K., et al., Depletion of adipocyte sphingosine kinase 1 leads to cell hypertrophy, impaired lipolysis, and nonalcoholic fatty liver disease. J Lipid Res, 2020. 61(10): p. 1328–1340.

7. Chaurasia, B., et al., Adipocyte Ceramides Regulate Subcutaneous Adipose Browning, Inflammation, and Metabolism. Cell Metab, 2016. 24(6): p. 820–834.

8. Chaurasia, B., et al., Ceramides are necessary and sufficient for diet-induced impairment of thermogenic adipocytes. Mol Metab, 2021. 45: p. 101145.

9. Gohlke, S., et al., Identification of functional lipid metabolism biomarkers of brown adipose tissue aging. Molecular metabolism, 2019. 24: p. 1–17.

10. Schweizer, S., et al., The lipidome of primary murine white, brite, and brown adipocytes-Impact of beta-adrenergic stimulation. PLoS Biol, 2019. 17(8): p. e3000412.

11. Coleman, D.L., Thermogenesis in diabetes-obesity syndromes in mutant mice. Diabetologia, 1982. 22(3): p. 205–11.

12. Saito, M., et al., Brown Adipose Tissue, Diet-Induced Thermogenesis, and Thermogenic Food Ingredients: From Mice to Men. Front Endocrinol (Lausanne), 2020. 11: p. 222.

13. Umekawa, T., et al., Effect of CL316,243, a highly specific beta(3)-adrenoceptor agonist, on lipolysis of epididymal, mesenteric and subcutaneous adipocytes in rats. Endocr J, 1997. 44(1): p. 181–5.

14. Umekawa, T., et al., Anti-obesity and anti-diabetic effects of CL316,243, a highly specific beta 3-adrenoceptor agonist, in Otsuka Long-Evans Tokushima Fatty rats: induction of uncoupling protein and activation of glucose transporter 4 in white fat. Eur J Endocrinol, 1997. 136(4): p. 429–37.

15. Yoneshiro, T., et al., Recruited brown adipose tissue as an antiobesity agent in humans. J Clin Invest, 2013. 123(8): p. 3404–8.

16. Morishige, J.-i., et al., Sphingosine kinase 1 is involved in triglyceride breakdown by maintaining lysosomal integrity in brown adipocytes. Journal of Lipid Research, 2023. 64(11).

17. Zhang, W., et al., Adipocyte lipolysis-stimulated interleukin-6 production requires sphingosine kinase 1 activity. Journal of Biological Chemistry, 2014. 289(46): p. 32178–32185.

18. Zhao, R.Z., et al., Mitochondrial electron transport chain, ROS generation and uncoupling (Review). Int J Mol Med, 2019. 44(1): p. 3–15.

19. Miller, C.N., et al., Isoproterenol Increases Uncoupling, Glycolysis, and Markers of Beiging in Mature 3T3-L1 Adipocytes. PLoS One, 2015. 10(9): p. e0138344.

20. Vijgen, G.H., et al., Increased oxygen consumption in human adipose tissue from the “brown adipose tissue” region. J Clin Endocrinol Metab, 2013. 98(7): p. E1230–4.

21. Acin-Perez, R., et al., A novel approach to measure mitochondrial respiration in frozen biological samples. EMBO J, 2020. 39(13): p. e104073.

22. Osto, C., et al., Measuring Mitochondrial Respiration in Previously Frozen Biological Samples. Curr Protoc Cell Biol, 2020. 89(1): p. e116.

23. Bridge-Comer, P.E. and S.M. Reilly, Measuring the Rate of Lipolysis in Ex vivo Murine Adipose Tissue and Primary Preadipocytes Differentiated In Vitro. J Vis Exp, 2023(193).

24. Hong, S., et al., Phosphorylation of Beta-3 adrenergic receptor at serine 247 by ERK MAP kinase drives lipolysis in obese adipocytes. Mol Metab, 2018. 12: p. 25–38.

25. Roy, D., J.M. Myers, and A. Tedeschi, Protocol for assessing ex vivo lipolysis of murine adipose tissue. STAR Protoc, 2022. 3(3): p. 101518.

26. de-Lima-Junior, J.C., et al., Abnormal brown adipose tissue mitochondrial structure and function in IL10 deficiency. EBioMedicine, 2019. 39: p. 436–447.

27. Doherty, E. and A. Perl, Measurement of Mitochondrial Mass by Flow Cytometry during Oxidative Stress. React Oxyg Species (Apex), 2017. 4(10): p. 275–283.

28. Pickles, S., N. Arbour, and C. Vande Velde, Immunodetection of outer membrane proteins by flow cytometry of isolated mitochondria. J Vis Exp, 2014(91): p. 51887.

